# Species-Specific gB Ectodomain Interactions and Cytoplasmic Domain Stability Regulate Herpes Simplex Virus Fusion

**DOI:** 10.1101/2025.10.25.684586

**Authors:** Qing Fan, Richard Longnecker, Sarah A. Connolly

## Abstract

Entry of herpesviruses into cells requires coordinated action of multiple viral glycoproteins, including gH/gL and gB which comprise the core fusion machinery conserved in herpesviruses. The gH/gL heterodimer activates the gB fusion protein, triggering its refolding from a prefusion to a postfusion form to drive membrane merger. The cytoplasmic tail domain (CTD) of gB is proposed to act as an inhibitory clamp that stabilizes the prefusion state, with interactions between gH and gB CTDs destabilizing this clamp. We previously found that herpes simplex virus 1 (HSV-1) and saimiriine herpesvirus 1 (SaHV-1) gB homologs are functionally interchangeable but mediate reduced fusion when coexpressed with heterotypic gH/gL. To map the regions of gB responsible for species-specific interactions, we generated HSV-1/SaHV-1 gB chimeras by swapping ectodomain, membrane-proximal region (MPR), transmembrane domain (TMD), and CTD segments. Our results show that homotypic CTD interactions alone are insufficient to trigger fusion, suggesting that gH/gL contacts the ectodomain of gB. We show that the HSV-1 gB CTD is hyperfusogenic relative to the SaHV-1 CTD, whereas the HSV-1 MPR is hypofusogenic relative to the SaHV-1 MPR. Together, these findings suggest that functional interaction between gH/gL-gB occur on both sides of the membrane and that gB maintains a balance between promotion and restraint of fusion through coordinated contributions of its domains.

## IMPORTANCE

HSV-1 entry requires the coordinated interaction of gD, gH/gL, and gB. Both gH/gL and gB are conserved herpesvirus proteins that are required for viral replication and are key targets of neutralizing antibodies. Despite their importance, how these proteins interact to mediate herpesvirus entry into cells remains poorly understood. In this study, we examined gB function by creating chimeras that swapped distinct domains between HSV-1 and SaHV-1 homologs. Using these chimeras, we demonstrate that a species-specific interaction occurs in the gB ectodomain, suggesting that gH/gL interacts with both the gB ectodomain and cytoplasmic tail. Additionally, we found that the HSV-1 CTD is hyperfusogenic compared to SaHV-1, suggesting that different gB domains can compensate for one another to balance fusion. This study provides new insight into how gB is regulated to mediate virus entry at the right time and place.

## INTRODUCTION

Herpesviruses are ubiquitous in humans, and the nine human herpesviruses cause diseases ranging from the common cold sore to life-threatening lymphomas. Herpes simplex virus type 1 (HSV-1) is an alphaherpesvirus that often manifests as recurrent mucocutaneous lesions localized to the face, mouth, cornea, or genitalia and, in rare instances, can lead to meningitis or encephalitis. Lesions can recur because HSV establishes a lifelong latent infection in sensory ganglia.

Herpesviruses are enveloped viruses that infect host cells by fusing the viral envelope with the host cell membrane. Their entry into cells requires coordinated interactions among several viral proteins. All nine human herpesviruses use a similar set of protein interactions to enter cells; thus, understanding the HSV-1 entry mechanism will provide insight into a conserved herpesvirus entry mechanism and inform the development of new therapies and prophylaxis.

Entry of HSV-1 into cells requires four viral glycoproteins that are necessary and sufficient for fusion: gD, gH, gL, and gB (Fig. 1A). The current model of entry places gH/gL between gD and gB in the sequence of events, with receptor binding by gD signaling a change in gH/gL that then triggers gB to refold and drive fusion (1–6). Several receptors for gD exist, including nectin-1, HVEM, and 3-O-sulfonated heparan sulfate (3-OS-HS). While HVEM is specific for HSV entry, nectin-1 is used as a receptor by a wide range of alphaherpesviruses, and related proteins such as CD155 can mediate fusion of other alphaherpesviruses. Additionally, receptors that bind HSV-1 gB have been reported to mediate fusion (7–9).

**Figure 1.**
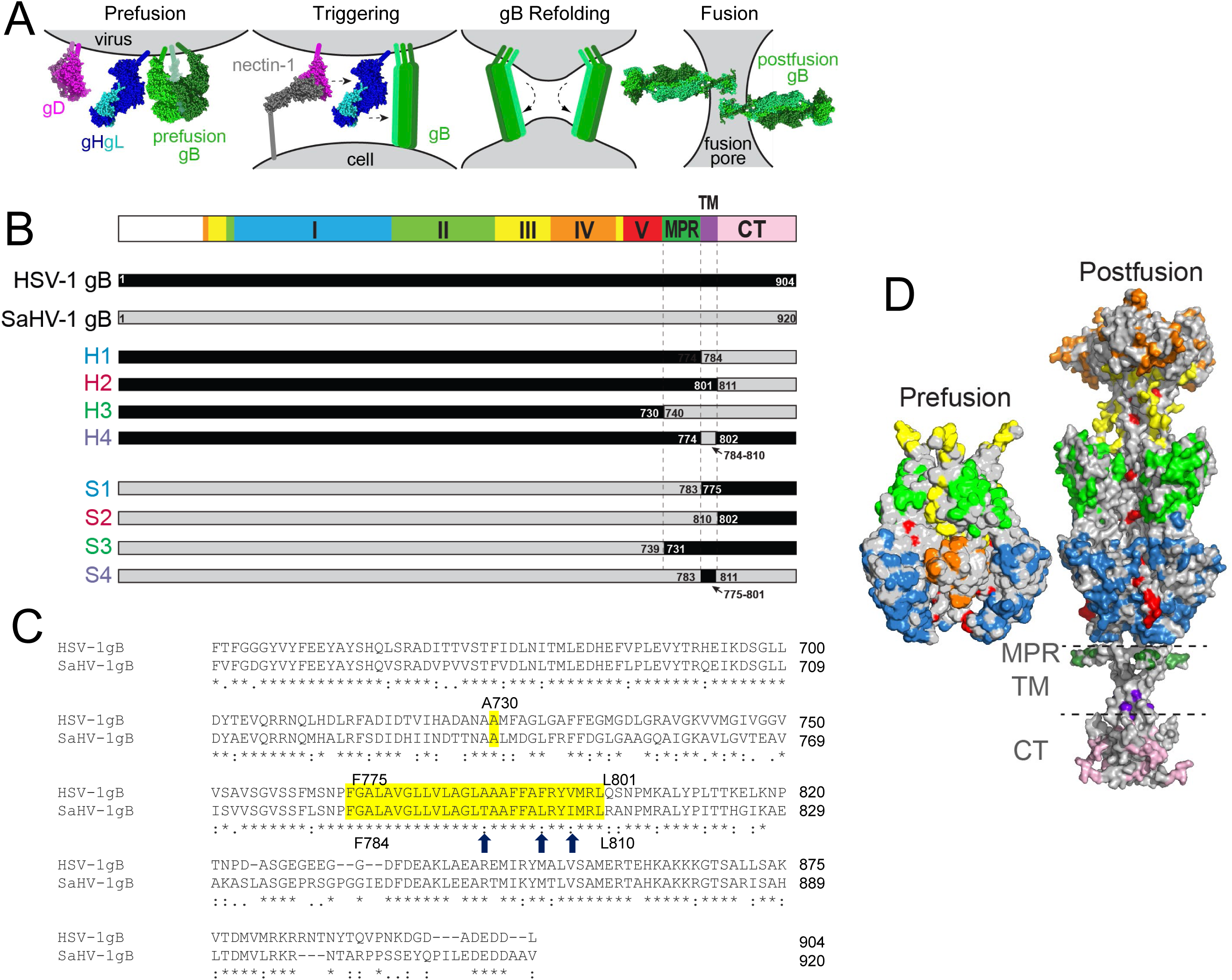
gB chimera constructs. (A) Model of HSV-1 entry into cells. gD dimer binds to nectin-1 and interacts with gH/gL heterodimer. gH/gL then signals gB trimer to refold from a prefusion to a postfusion conformation. The refolding of multiple gB trimers drives fusion of the viral and cellular membranes. (B) Schematic representation of chimeric gB constructs comprised of HSV-1 (black) and SaHV-1 (gray) sequences. The top bar illustrates the four regions of gB that are swapped: the N-terminal ectodomain (domains I-V in blue, green, yellow, orange, and red), the membrane-proximal region (MPR, dark green), the transmembrane region (TMD, purple), and the cytoplasmic tail (CTD, pink). Reciprocal constructs with N-terminal ectodomain from HSV-1 (H1-H4) or SaHV-1 (S1-S4) are shown. Residues that serve as break points for the chimeras are shown. (C) Sequence alignment of the C-terminal regions from HSV-1 and SaHV-1 gB. HSV-1 residue 730 (yellow) marks the end of the N-terminal region of the ectodomain, prior to the MPR. The TMD (yellow) spans HSV-1 gB residues F775-L801 and SaHV-1 gB residues F784-L810. Only three of the TMD residues differ between species (arrows). (D) Surface rendering of prefusion gB cryoelectron microscopy model (PDB 6Z9M) and postfusion gB crystal structure (PDB 5V2S). Only the ectodomain was modeled for prefusion gB, whereas the full-length model of postfusion gB is shown, anchored in a membrane (dashed lines). Residues that differ between HSV-1 and SaHV-1 gB are colored by domain (DI blue, DII green, DIII yellow, DIV orange, DV red, MPR dark green, TM purple, CT pink).

gB is a class III viral fusion protein that is conserved across all herpesviruses and structurally homologous to the fusion proteins of rhabdoviruses, thogotoviruses, and baculoviruses (10–12). Viral fusion proteins are metastable proteins that insert into a target membrane and refold from a prefusion to a postfusion conformation to bring the viral and cell membranes together (Fig. 1A). The prefusion and postfusion structures of gB have been solved for multiple herpesviruses(1, 13–16). HSV-1 gB is a trimer consisting of eight domains (Fig. 1B). Domain I (DI) includes hydrophobic fusion loops that insert into the cell membrane. DIII forms a trimeric coiled coil at the core of the postfusion structure, against which a domain V arm packs in an anti-parallel orientation. The membrane proximal region (MPR), transmembrane domain (TMD) and cytoplasmic tail domain (CTD) form a trimeric pedestal that anchors the gB ectodomain in the membrane (Fig. 1D) (17).

Chimeric constructs from homologous herpesviruses have been used to investigate homotypic interactions among entry glycoproteins. We previously examined homotypic requirements for entry glycoproteins from HSV-1 and saimiriine herpesvirus 1 (SaHV-1), a primate alphaherpesvirus with sequence similarity with HSV-1(18, 19). We found that HSV-1 and SaHV-1 gD and gH/gL function in fusion in a species-specific manner, and we leveraged that species-specificity to map sites of gD-gH/gL functional interaction sites by assessing the fusion function of chimeric proteins carrying HSV-1 and SaHV-1 segments.

Unlike gD and gH/gL, HSV-1 and SaHV-1 gB homologs can be functionally swapped in a cell-cell fusion assay (18). In fact, SaHV-1 gB supports low levels of virus entry when added to the HSV genome (20). Although these results demonstrate that gB retains function with heterotypic gD and gH/gH, gB proteins function best when coexpressed with homotypic entry glycoproteins. To identify the regions of gB that impart this homotypic species preference for function, the current study generated a panel of gB chimeras carrying HSV-1 and SaHV-1 segments by swapping the ectodomains, membrane-proximal regions (MPR), transmembrane domains (TM) and/or cytoplasmic tails (CT) (Fig. 1B).

## MATERIALS AND METHODS

### Cells and antibodies

Chinese hamster ovary (CHO-K1) (ATCC, USA) cells were grown in Ham’s F12 medium supplemented with 10% FBS. C10 cells, derived from B78-H1 mouse melanoma cell line to stably expresses human nectin-1 (21), were grown in DMEM supplemented with 10% FBS and 500 μg/mL of G418. Anti-HSV-1 gB polyclonal antibody (PAb) R74 (22), recognizes HSV-1 gB and SaHV-1 gB. Anti-Flag monoclonal antibody (Sigma, F1804) recognizes Flag-tagged gB.

### Plasmids

Plasmids expressing HSV-1(KOS) gB (pPEP98), gD (pPEP99), gH (pPEP100) and gL (pPEP101) were previously described (23), as were plasmids expressing human nectin-1 (pBG38) (24), human HVEM (pBEC10) (25) and CD155 (kindly provided by Dr. Eckard Wimmer). Plasmids expressing SaHV-1 gB (pQF77), gD (pQF78), gH (pQF79) and gL (pQF80) were previously described (18), as well as Flag-tagged HSV-1 gB (pQF112) and SaHV-1 gB (pQF81) (18).

Fig. 1B depicts the constructs generated for this study. All the new constructs were cloned into pCAGGS using PCR products amplified from pPEP98 or pQF77. H1 (pQF494) includes M1-P774 of HSV-1 gB and F784-V920 of SaHV-1 gB. H2 (pQF495) includes M1-L801of HSV-1 gB and R811-V920 of SaHV-1 gB. H3 (pQF493) includes M1-A730 of HSV-1 gB and L740-V920 of SaHV-1 gB. For H4 (pQF498), the TMR of HSV-1 gB (F775-L801) was replaced with the TMR of SaHV-1 gB (F784-L810). S1 (pQF489) includes M1-P783 of SaHV-1 gB and F775-L904 of HSV-1 gB. S2 (pQF491) includes M1-L810 of SaHV-1 gB and Q802-L904 of HSV-1 gB. S3 (pQF487) includes M1-A739 of SaHV-1 gB and M731-L904 of HSV-1 gB. For S4 (pQF497), the TMR of SaHV-1 gB (F784-L810) was replaced with the TMR of HSV-1 gB (F775-L801). All plasmids generated for this study were sequenced by ACGT, Inc. (Wheeling, IL 60090, USA).

### CELISA

Cell-based ELISA (CELISA) was used to evaluate the cell surface expression of the HSV-1 and SaHV-1 gB chimera proteins. CHO-K1 cells seeded in 96-well plates were transfected with 60 ng/well of empty vector or plasmids expressing HSV-1 gB, SaHV-1 gB or the chimeras using 0.15 µl of Lipofectamine 2000 (Invitrogen) in Opti-MEM (Invitrogen). The cells were washed once with PBS 24 h after transfection, and CELISA was performed as described (26). Anti-gB R74 polyclonal antibody was used to probe for expression of the gB constructs, and anti-Flag monoclonal antibody (Sigma F1084) was used to compare expression of the Flag-tagged WT gB constructs. After incubation with primary antibody, cells were washed, fixed, and incubated with biotinylated goat anti-rabbit IgG (Sigma) for gB constructs or biotinylated goat anti-mouse IgG (Sigma) for the Flag-tagged constructs, followed by streptavidin-HRP (GE Healthcare) and HRP substrate (BioFX).

### Cell-cell fusion assay

Fusion activity was measured using a quantitative luciferase-based cell fusion assay as previously described (23). CHO-K1 cells were seeded in 6-well plates overnight. One set of CHO-K1 cells (effector cells) was transfected with 2 µg DNA/well, including 400 ng each of plasmids encoding T7 RNA polymerase, a version of gB (WT, chimera, or vector), and gD, gH, and gL from HSV-1 or SaHV-1, using 5 µl of Lipofectamine 2000. A second set of CHO-K1 cells (target cells) was transfected with 400 ng/well of a plasmid encoding the firefly luciferase gene under control of the T7 promoter plus 1.5 µg/well of empty vector (pcDNA3) or plasmid encoding receptors (nectin-1, HVEM, or CD155) using 5 µL of Lipofectamine 2000. Six h post-transfection, the cells were detached with versene and suspended in 1.5 ml of F12 medium supplemented with 10% FBS. Effector and target cells were mixed in a 1:1 ratio and replated in 96-well plates for 18 h. Luciferase activity was quantitated using a luciferase reporter assay system (Promega) and a Wallac-Victor luminometer (Perkin Elmer).

### Syncytium formation assay

The syncytium formation assay was performed as described previously (27, 28). C10 cells in 24-well plates were transfected with 1 µg/well DNA, including 250 ng each of plasmids encoding gB (WT, chimera, or vector), gD, gH, and gL, using 2.5 µL FuGENE 6 (Promega, USA) in 500 µL DMEM supplemented with 10% FBS. After overnight incubation, the transfected cells were washed with PBS, fixed with methanol, and stained with Giemsa for 30 minutes. Cells were imaged using an EVOS Cell Imaging System. Syncytia, defined as multinucleated cells containing three or more nuclei, were identified by microscopy, and nuclei within each syncytium were counted either directly under the microscope or from captured images.

## RESULTS

### Design of gB chimeras

Our previous study demonstrated that HSV-1 gB and SaHV-1 gB could be swapped in a fusion assay, indicating that the gB retains function when coexpressed with gD and gH/gL from the heterotypic species (18). However, the gB proteins function best when coexpressed with entry glycoproteins partners from their homotypic species. In the previous study, SaHV-1 gB retained 15% of its fusion function when coexpressed with HSV-1 gD and gH/gL, and HSV-1 gB retained 66% of fusion function when coexpressed with SaHV-1 gD and gH/gL. These deficiencies in fusion indicated that species-specific interactions contribute to triggering gB during the fusion event.

HSV-1 and SaHV-1 gB share 65% sequence identity (Fig. 1D). To identify the regions of gB that impart species-specificity to fusion, the residues comprising the ectodomain, MPR, TMD and CTD were identified by sequence alignment and chimeric versions of gB including combinations of these regions from either species were constructed (Fig. 1B).

### Expression of gB chimeras

To assess whether HSV-1 and SaHV-1 gB typically are expressed at equivalent levels, the cell surface expression of N-terminally Flag-tagged versions of HSV-1 gB and SaHV-1 gB was examined in transfected CHO-K1 cells by CELISA using an anti-Flag MAb (Fig. 2A) (26, 29). Both Flag-tagged gB constructs were expressed at similar levels.

**Figure 2.**
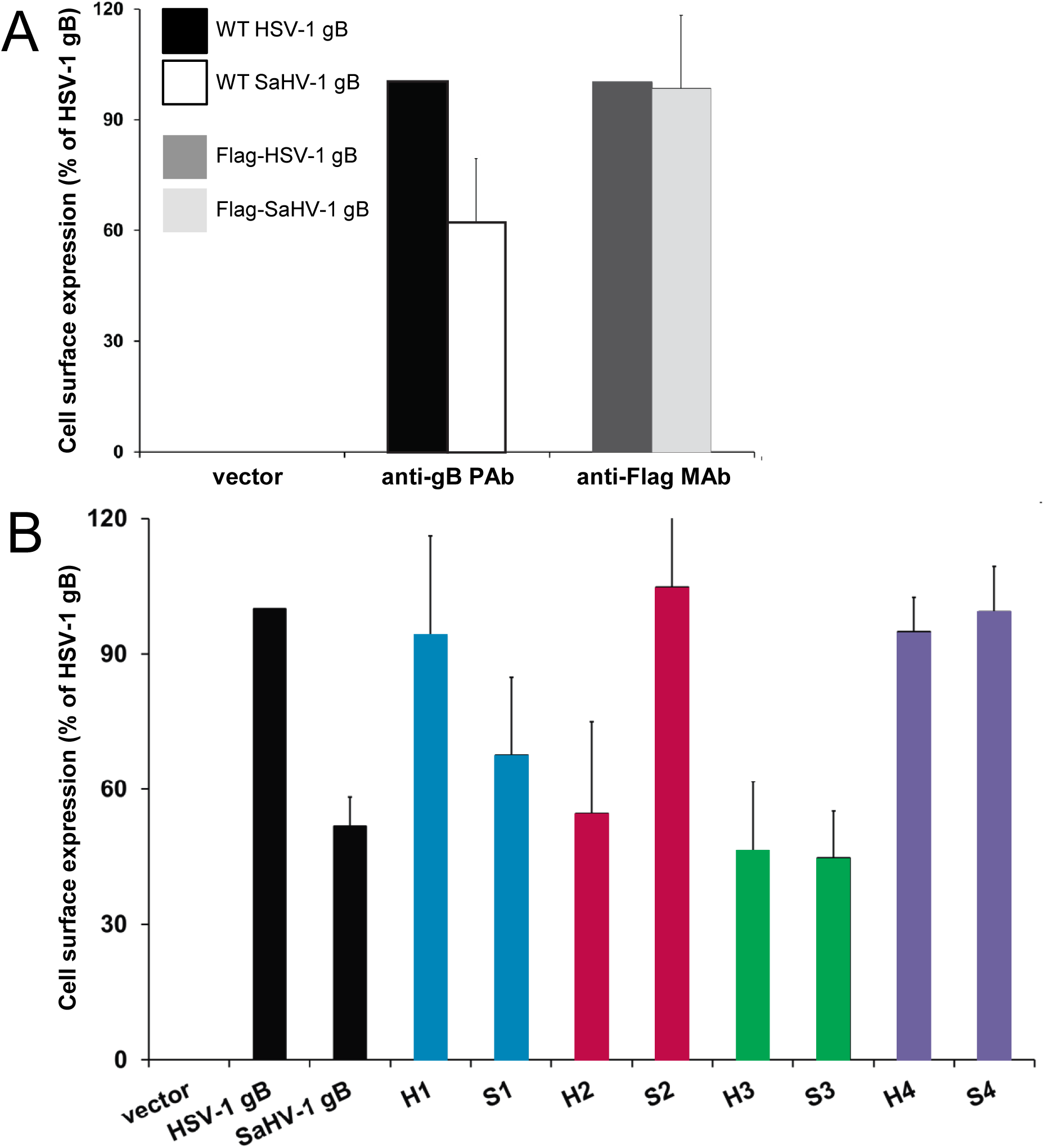
Cell surface expression of HSV-1 and SaHV-1 gB. (A) CHO-K1 cells were transfected overnight with plasmids encoding WT or Flag-tagged HSV-1 or SaHV-1 gB or empty vector. Cell surface gB expression was detected by adding a polyclonal anti-HSV-1 gB antibody (R74) or a monoclonal anti-Flag antibody (F1804 M2), followed by fixative and secondary antibody. (B) CHO-K1 cells were transfected overnight with plasmids encoding WT gB, gB chimeras, or an empty vector. Cell surface gB expression was detected by adding a R74 PAb, followed by fixative and secondary antibody. Each bar shows the mean and standard deviation of three independent experiments. Background signals from the vector only control were subtracted from the values. Data are normalized to the expression level of HSV-1 gB.

To determine how well the anti-HSV-1 gB polyclonal antibody (PAb) R74 would cross-react with SaHV-1 gB, cell surface expression of WT forms of gB was detected by CELISA using R74. WT SaHV-1 gB was detected at the cell surface at approximately 50% of the level of WT HSV-1 gB. Despite this reduction, this level of R74 reactivity was sufficient to assess whether the chimeric constructs reached the cell surface without the need to add a Flag-tag to the constructs.

Cell surface expression of the gB chimeras was evaluated by CELISA using R74 (Fig. 2B). All constructs were detected at the cell surface, suggesting appropriate trafficking of the gB with proper conformation. Constructs including the HSV-1 gB ectodomain (H1-H4) were expressed at 45-95% of the level of WT HSV-1 gB. Constructs including the SaHV-1 gB ectodomain (S1-S4) showed surface expression at 45-105%, compared to HSV-1 gB. These CELISA results show that, although some of the constructs have reduced cell surface expression, all constructs were expressed on the cell surface at levels that permitted further evaluation for fusion function.

### Fusion function of gB chimeras

Our previous study showed that SaHV-1 glycoproteins mediated fusion nearly as well as HSV-1 glycoproteins with cells expressing nectin-1 (18). To determine the species-specificity of fusion displayed by each of the gB chimeras, we used a quantitative cell-cell fusion assay to assess fusion activity when the chimeras were expressed with entry glycoproteins from either HSV-1 or SaHV-1 (Fig. 3). CHO-K1 cells (target cells) were transfected with nectin-1 receptor and a plasmid encoding luciferase under a T7 promoter. A second set of CHO-K1 cells (effector cells) were transfected with a gB chimera plus gD, gH, and gL from either HSV-1 or SaHV-1 and T7 RNA polymerase. The effector and target cells were mixed and luciferase activity was recorded as a measure of cell-cell fusion.

**Figure 3.**
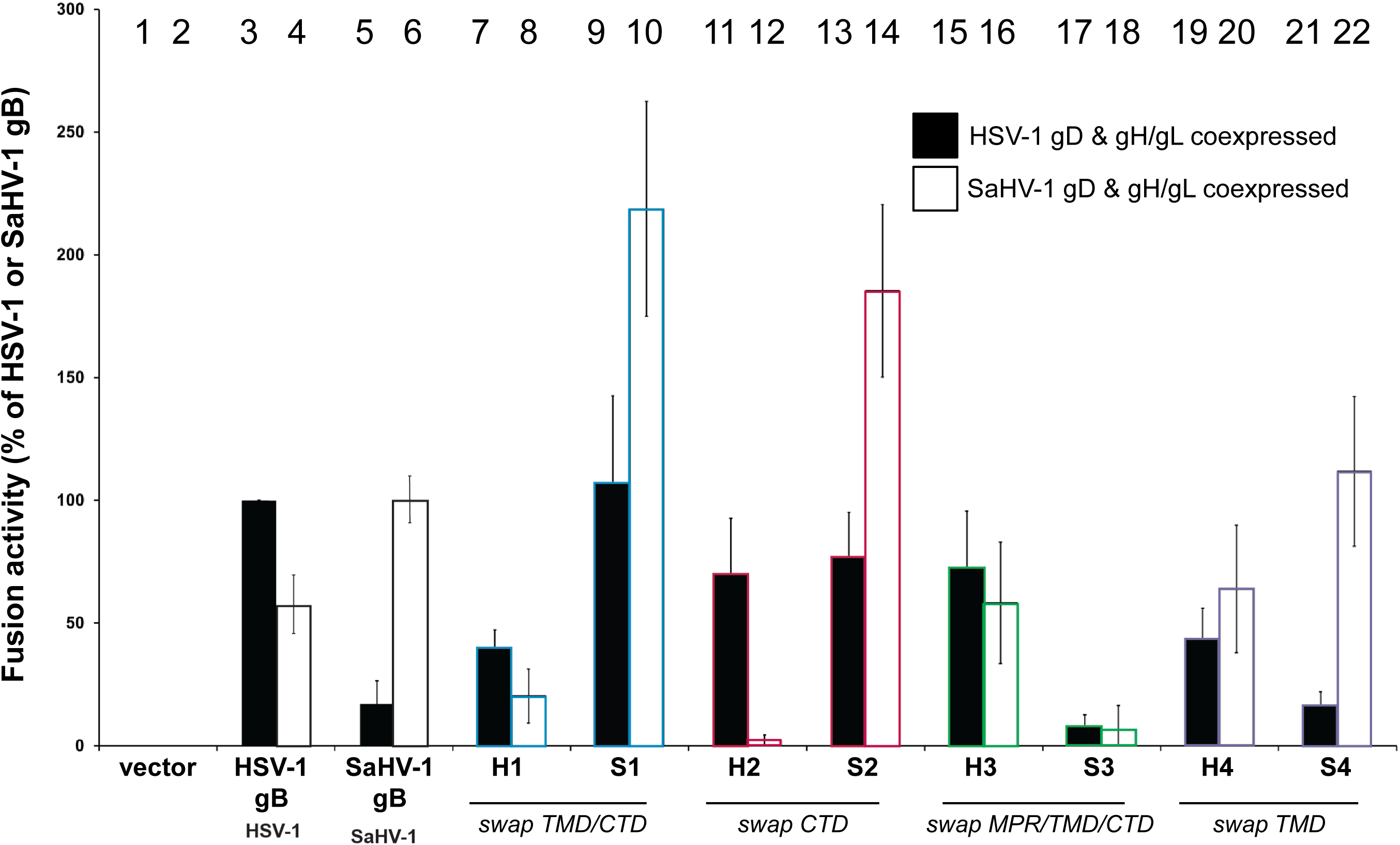
Cell-cell fusion activity of HSV-1 and SaHV-1 gB chimeras with nectin-1. One set of CHO-K1 cells (effector cells) was transfected with plasmids encoding a version of gB (WT, chimera, or empty vector), gD, gH, and gL from either HSV-1 (filled bars) or SaHV-1 (open bars), and T7 polymerase. A second set of CHO-K1 cells (target cells) was transfected with plasmids carrying the luciferase gene under the control of the T7 promoter and nectin-1 receptor. Target and effector cells were cocultured for 18 h and luciferase activity was measured as an indication of cell-cell fusion. Data are expressed as a percentage of fusion mediated by WT HSV-1 gB for the HSV-1 gD and gH/gL samples (filled bars) or WT SaHV-1 gB for the SaHV-1 gD and gH/gL samples (open bars). Background fusion detected after transfection with vector instead of gB was subtracted from the values. Means and standard deviations of results of three independent experiments are shown. Bars are numbered for reference. The regions that were swapped are noted below the reciprocal chimera pairs.

### The gB ectodomain contributes to fusion in a species-specific manner

Previous studies using bimolecular fluorescence complementation suggested that the gB and gH/gL ectodomains interact (30, 31). To examine species-specific interactions of the gB ectodomain, the gB ectodomains were swapped in the H1 and S1 chimeras. The H1 chimera includes the ectodomain of HSV-1 with the TMD and CTD of SaHV-1, whereas the S1 chimera conversely includes the ectodomain from SaHV-1 with the TMD and CTD from HSV-1 (Fig. 1). H1 demonstrated higher levels of fusion when coexpressed with HSV-1 gD and gH/gL, as compared to its fusion when coexpressed with SaHV-1 gD and gH/gL (Fig. 3, bars 7 and 8). Likewise, S1 demonstrated higher levels of fusion when coexpressed with SaHV-1 gD and gH/gL, rather than HSV-1 gD and gH/gL (Fig. 3, bars 9 and 10). These results indicate that a species-specific fusion element in gB maps to the ectodomain and suggest that the gB ectodomain interacts with other glycoproteins, presumably gH/gL, in a species-specific manner.

Compared to WT SaHV-1 gB (Fig. 3, bars 5 and 6), S1 fusion was increased markedly when coexpressed with either HSV-1 or SaHV-1 entry glycoproteins (Fig. 3, bars 9 and 10). Conversely, H1 fusion was decreased compared to WT HSV-1 gB (Fig. 3, bars 3 and 4) when coexpressed with HSV-1 entry glycoproteins and unchanged when coexpressed with SaHV-1 entry glycoproteins. Since the enhanced fusion of S1 was maintained when it was coexpressed with glycoprotein partners from either species, these results suggest that the SaHV-1 gB ectodomain is more fusogenic overall than the ectodomain of HSV-1 gB.

### Interactions with the gB cytoplasmic tail are insufficient to trigger fusion

Mutations in the gB CTD can inhibit or enhance fusion (31–44) and an interaction between the CTD of gB and gH has been proposed (45). To explore species-specific requirements for the gB CT, gB CTD was swapped in the H2 and S2 chimeras. H2 includes the ectodomain and TMD of HSV-1 with the CTD of SaHV-1, whereas S2 includes the ectodomain and TMD from SaHV-1 with the CTD of HSV-1 (Fig. 1). H2 mediates fusion well when coexpressed with HSV-1 gD and gH/gL, but it fails to mediate fusion when coexpressed with SaHV-1 gD and gH/gL (Fig. 3, bars 11 and 12). These results indicate that a species-specific element for fusion promotion is present in the ectodomain and/or TM, consistent with the previous result for H1. Similarly, consistent with the results for S1, S2 mediates fusion better when coexpressed with SaHV-1 gD and gH/rather than HSV-1 gD and gH/gL (Fig. 3, bars 13 and 14), suggesting a species-specific interaction with the ectodomain and/or TM.

These H2 and S2 results show that the gB CTD alone is insufficient for interacting with the entry glycoprotein partners that trigger fusion. H2 carries the SaHV-1 gB CTD but does not mediate fusion when coexpressed with gD and gH/gL from SaHV-1 (Fig. 3 bar 12). The fact that H2 mediates fusion when expressed with HSV-1 glycoproteins indicates that H2 is expressed and competent for fusion. This result suggests that an interaction between SaHV-1 gB CTD and gH/gL is insufficient to trigger fusion. Alternatively, the H2 CTD conformation could be altered compared to that of WT SaHV-1 gB.

### The HSV-1 gB cytoplasmic tail is hyperfusogenic compared to the SaHV1 gB CT

As observed for S1, S2 fusion was increased compared to WT SaHV-1 gB, when coexpressed with gD and gH/gL from either HSV-1 or SaHV-1 (Fig. 3, bars 5, 6, 13, and 14). Correspondingly, H2 fusion was decreased when coexpressed with gD and gH/gL from either HSV-1 or SaHV-1 (Fig. 3, bars 3, 4, 11, and 12). These results demonstrate a global effect of the gB CTD on fusion. These results suggest the CTD of HSV-1 gB is hyperfusogenic when compared to the CTD of SaHV-1 gB. The HSV-1 CTD may be less stable than that of SaHV-1 and/or the ectodomain of SaHV-1 gB may be less stable than that of HSV-1. The result is clearer when comparing S2 and WT SaHV-1 gB fusion (Fig. 3, bar 14) when coexpressed with SaHV-1 gD and gH/gL (Fig. 3, bar 6). In summary, fusion mediated by SaHV-1 gB was enhanced by the addition of the HSV-1 gB CTD (i.e. S2), whereas fusion mediated by HSV-1 gB was impaired by the addition of the SaHV-1 gB CTD (i.e. H2).

### The SaHV-1 gB MPR is hyperfusogenic compared to the HSV-1 gB MPR

The MPR comprises the C-terminal end of the ectodomain and includes a C-terminal helix that is embedded in and parallel to the outer leaflet of the membrane in the postfusion form of gB (17). Mutations in the MPR can abrogate infectivity (33). To evaluate a species-specific role of the MPR, chimeras were generated with a swap point at the start of the MPR. H3 includes most of the HSV-1 gB ectodomain with SaHV-1 residues for the MPR, TM, and CTD (Fig. 1). When coexpressed with either HSV-1 or SaHV-1 gD and gH/gL, H3 mediates fusion better than H1 (Fig. 3, bars 7, 8, 15, and 16). These results indicate that the SaHV-1 MPR present in H3 enhances fusion in both an HSV-1 and SaHV-1 background.

The complementary construct S3 includes most of the SaHV-1 gB ectodomain with the MPR, TM, and CTD from HSV-1. When coexpressed with either HSV-1 or SaHV-1 gD and gH/gL, S3 shows less fusion than S1 (Fig. 3, bars 13, 14, 17, and 18). In agreement with the H3 data interpretation, these S3 results suggest that the HSV-1 MPR dampens fusion in both an HSV-1 and SaHV-1 background. Alternatively, although S3 expresses on the cell surface at nearly the same level as WT SaHV-1 gB, the fact S3 fails to mediate fusion well in both an HSV-1 and SaHV-1 entry glycoproteins could indicate that S3 is not properly folded.

These data demonstrate a global effect of the gB MPR on fusion and they do not support or refute a species-specific role of the gB MPR. The MPR of SaHV-1 gB may be destabilized compared to the HSV-1 gB MPR. An alignment of the gB MPR shows greater divergence towards the N-terminus of the MPR (Fig. 1C), a region that was not resolved in the crystal structure (17).

### The gB TMD does not contribute to the species-specificity of fusion

Mutations in the gB TMD also have been shown to inhibit infectivity (33). An alignment of the HSV-1 and SaHV-1 TMD domains shows only three residue differences (Fig. 1C). To determine if these TMD residues promote fusion in a species-specific manner, chimeras were generated with the only TMD domains exchanged. H4 includes HSV-1 gB sequence with a TMD from SaHV-1 gB, and S4 includes SaHV-1 gB sequence with a TMD from HSV-1 gB. S4 closely mirrored the fusion activity of SaHV-1 gB, mediating reduced fusion when coexpressed with HSV-1 gD and gH/gL as compared to SaHV-1 gD and gH/gL (Fig. 3, bars 21 and 22). Thus, the addition of the heterotypic HSV-1 gB TMD to S4 failed to enhance functional interaction with HSV-1 gD and gH/gL. Similarly, the addition of the SaHV-1 gB TMD to H4 did not enhance a functional interaction with SaHV-1 gD and gH/gL (Fig. 3, bars 4 and 20). H4 fusion was reduced compared to WT HSV-1 gB when co-expressed with HSV-1 gD and gH/gL (Fig. 3, bars 3 and 19), suggesting a possible modest role for the TMD in functional interactions or gB folding.

### The HSV-1 gB CTD remains hyperfusogenic when fusion is mediated by CD155 or HVEM

Whereas nectin-1 mediates entry of both HSV-1 and SaHV1 (24), HVEM can mediate fusion of HSV-1 but not SaHV-1, and conversely CD155 can mediate fusion of SaHV-1 but not HSV-1 (18). These restrictions most likely correlate with the ability of each gD homolog to bind the receptors. To examine whether gB chimera function was impacted by the gD receptor, fusion mediated by the chimeras was assessed when CD155 or HVEM were transfected in place of nectin-1 (Fig. 4).

**Figure 4.**
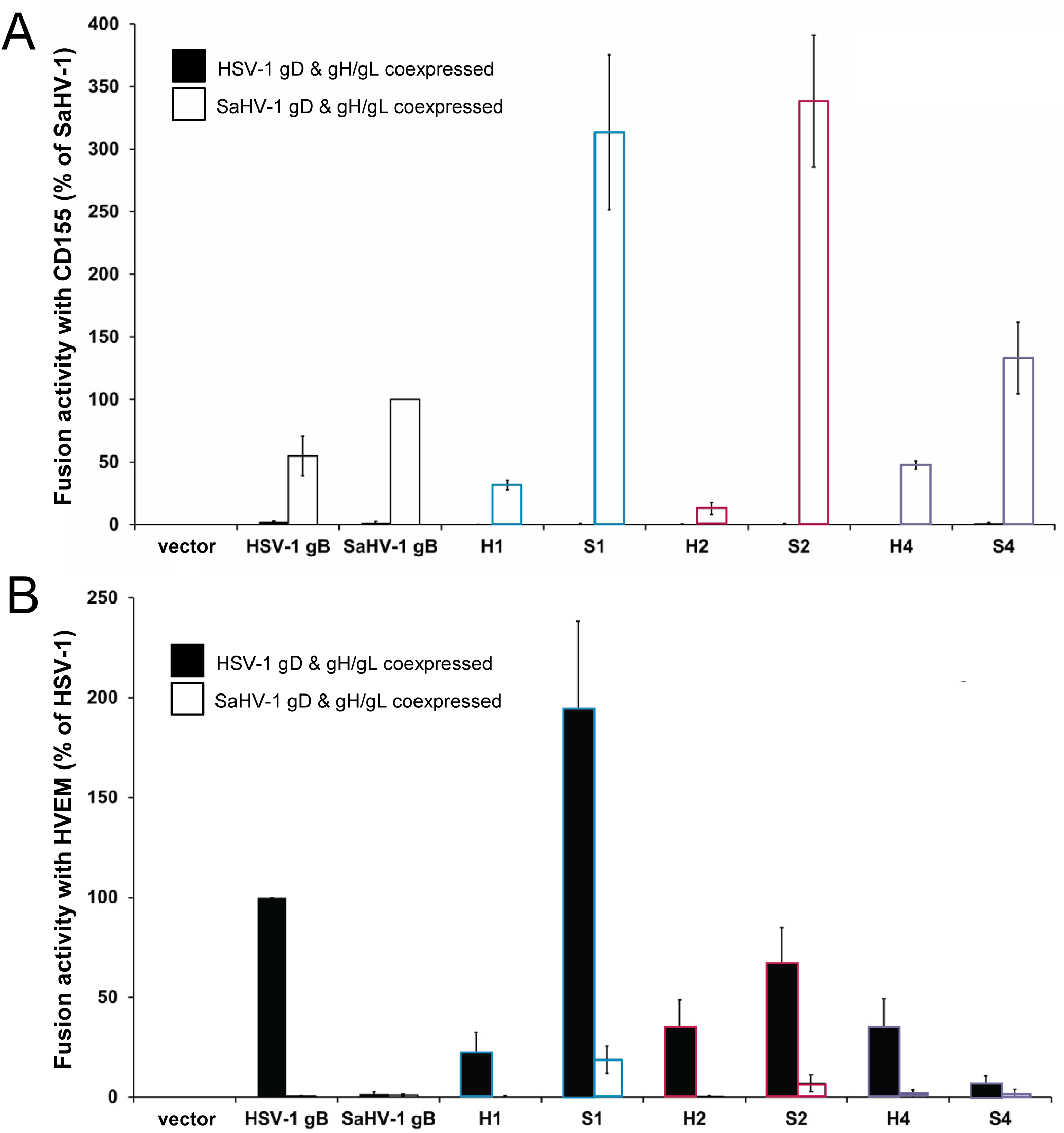
Cell-cell fusion activity of HSV-1 and SaHV-1 gB chimeras with CD155 or HVEM. Effector CHO-K1 cells were transfected with plasmids encoding a version of gB (WT, chimera, or empty vector), gD, gH, and gL from either HSV-1 (filled bars) or SaHV-1 (open bars), and T7 polymerase. Target cells were transfected with plasmids carrying the luciferase gene under the control of the T7 promoter and CD155 (A) or HVEM (B) receptor. Cells were cocultured and luciferase activity was measured. Data are shown as a percentage of WT SaHV-1 gB (for CD155) or WT HSV-1 gB (for HVEM), following subtraction of vector-only background signal. Means and standard deviations of results of three independent experiments are shown.

As reported previously, WT SaHV-1 gB coexpressed with SaHV-1 gD and gH/gL mediated fusion with CD155-expressing cells, but WT HSV-1 gB coexpressed with HSV-1 gD and gH/gL failed to mediate fusion (Fig. 4A). As expected, none of the gB constructs mediated fusion with CD155-expressing cells when coexpressed with HSV-1 gD and gH/gL because CD155 is not a receptor for HSV-1 gD. When coexpressed with SaHV-1 gD and gH/gL, S1 and S2 demonstrated enhanced fusion compared to SaHV-1 gB (Fig 4A), whereas H1 and H2 showed reduced fusion compared to HSV-1 gB. These results affirm the nectin-1 results indicating that the HSV-1 CTD is hyperfusogenic when added to the SaHV-1 gB ectodomain.

As expected, WT HSV-1 gB coexpressed with HSV-1 gD and gH/gL mediated fusion with HVEM-expressing cells, but WT SaHV-1 gB coexpressed with SaHV-1 gD and gH/gL failed to mediate fusion because HVEM is not a receptor for SaHV-1 gD (Fig. 4B). When coexpressed with HSV-1 gD and gH/gL, S1 and S2 displayed greatly enhanced fusion compared to SaHV-1 gB (Fig. 4B), confirming that the HSV-1 CTD is hyperfusogenic. In fact, both S1 and S2 mediated detectable levels of fusion even when coexpressed with SaHV-1 gD and gH/gL, despite the previous finding that SaHV-1 gD does not function with HVEM (18). Consistent with the nectin-1 results (Fig. 3), H1 and H2 showed reduced fusion compared to WT HSV-1 gB when coexpressed with HVEM and HSV gD and gH/gL, suggesting that the SaHV-1 CTD is hypofusogenic in comparison. Thus, the hyperfusogenic nature of the HSV-1 gB CTD was apparent when fusion was examined using all three gD receptors.

### S1 and S2 chimeras are hyperfusogenic in a syncytium formation assay

To evaluate S1 and S2 fusion using a second approach, we performed a syncytia assay with both constructs. The known hyperfusogenic mutant gB876t (3–5), which is truncated after CTD residue 876, was included as a control. B78-H1 cells expressing nectin-1 (C10 cells) (46) were transfected overnight with gD, gH/gL, and gB and syncytia were quantified (Fig. 4A). B78-H1 cells were selected because they do not express endogenous HSV-1 receptor(s), and they display a low level of background syncytium formation. S1 and S2 mediated enhanced levels of syncytia formation when coexpressed with SaHV-1 gD and gH/gL, consistent with the quantitative fusion assay (Fig. 5A). Unlike the quantitative fusion assay, S1 and S2 also showed enhanced syncytium formation when coexpressed with HSV-1 gD and gH/gL. Images of the syncytia show large multi-nucleated cells when S1 and S2 were coexpressed with either HSV-1 or SaHV-1 gD and gH/gL (Fig. 5B).

**Figure 5.**
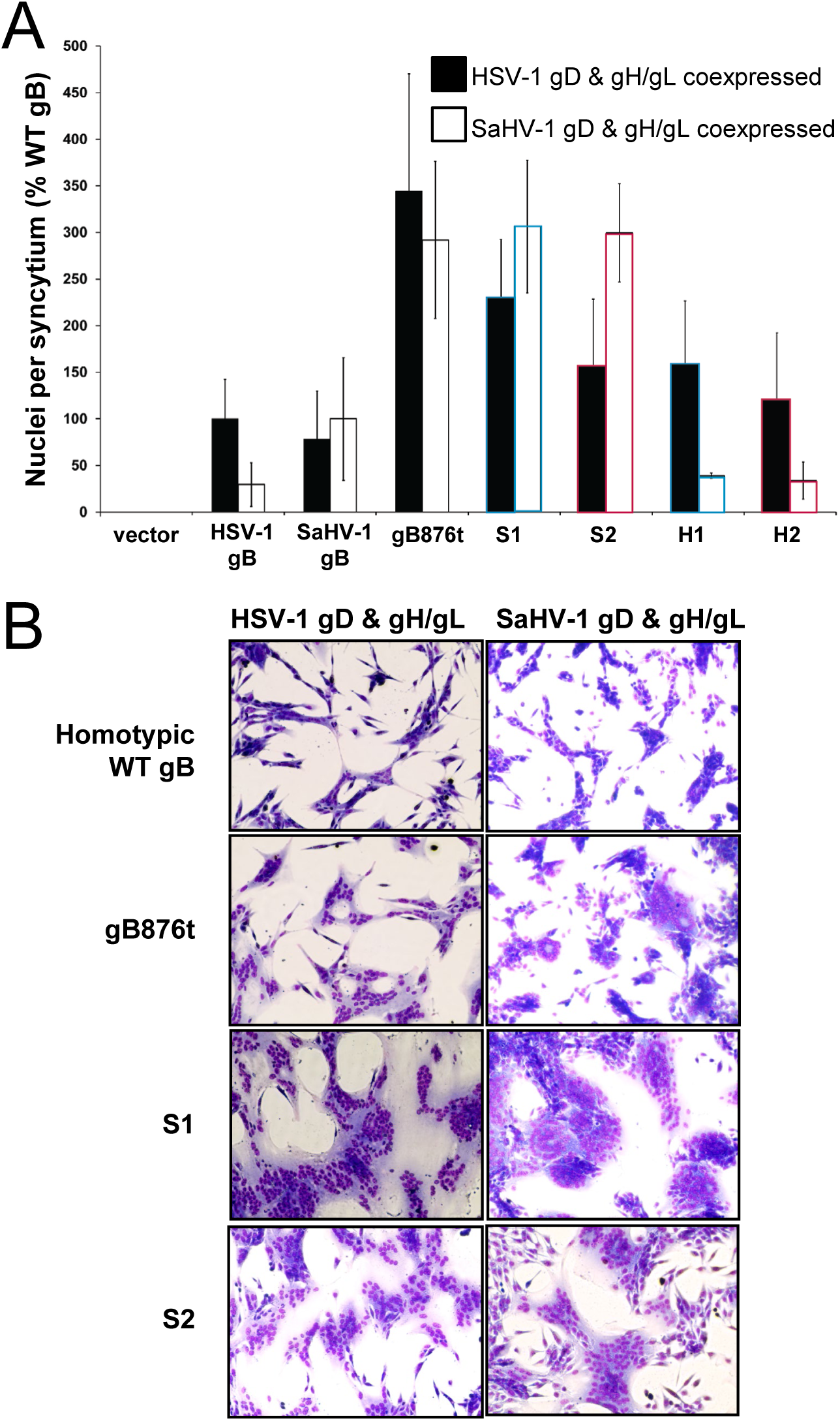
Syncytium formation by gB chimeras. (A) C10 cells stably expressing nectin-1 were transfected with plasmids encoding a version of gB (WT, chimera, or empty vector) and gD, gH, and gL from either HSV-1 (filled bars) or SaHV-1 (open bars). Cells were stained and imaged, and the number of nuclei per syncytium was counted. Data are presented as a percentage relative to the number of observed with WT glycoprotein expression. Results represent the means from at least three independent experiments. **(B)** Representative images show syncytia resulting from the fusion assays for WT gB and three hyperfusogenic mutants (gB876t, S1, and S2).

### S1 and S2 chimeras retain a requirement for gD and gH/gL co-expression to mediate fusion

Given the S1 and S2 hyperfusogenic phenotypes, we examined whether these chimeras were able to mediate fusion in the absence of signal from gD and gH/gL. The quantitative fusion assay was performed using nectin-1 expressing target cells, with gD excluded from some samples and gH/gL excluded from other samples (Fig. 5). Both S1 and S2 required coexpression of gD and gH/gL to mediate fusion, indicating that these hyperfusogenic mutants retained a canonical triggering requirement.

## DISCUSSION

Mapping the functional interaction sites on herpesvirus entry glycoprotein has been challenging because glycoprotein complex interactions are low affinity and/or transient. Surface plasmon resonance has shown that purified forms of the gD and gH/gL ectodomains interact directly (47, 48), and this interaction site has been validated using neutralizing antibodies (49). The interaction between gH/gL and gB is less well defined. An interaction between purified forms of the gH/gL and gB ectodomains was demonstrated using a liposome co-flotation assay (50). This association occurred only after treatment at low pH, which can trigger changes in gB conformation(6) and is relevant to endocytic route of entry(51, 52). Physical interactions between gB and gH/gL also have been shown using bimolecular fluorescence complementation (30, 53) and a split-luciferase assay (45); however, the functional relevance of these interactions is not clear from these approaches. The current study leverages the species-specificity of HSV-1 and SaHV-1 fusion to map functional interaction sites on gB using functional gB chimeras.

Our results indicate that the gB ectodomain contributes to fusion in a species-specific manner, presumably through a functional interaction with the gH/gL ectodomain. The data show that interactions with the gB CTD are insufficient to trigger fusion because fusion function is tracked with the homotypic gB ectodomain. Specifically, chimeras including the HSV-1 ectodomain (H1 and H2) demonstrated higher levels of fusion when coexpressed with HSV-1 gD and gH/gL (Fig. 3). Likewise, chimeras including the SaHV-1 ectodomain (S1 and S2) demonstrated higher levels of fusion when coexpressed with SaHV-1 gD and gH/gL, rather than HSV-1 gD and gH/gL. These results expand upon the previous liposome flotation assay finding that showed that the gH/gL and gB ectodomains can physically associate (50) because they demonstrate that the gB ectodomain interaction is functionally important.

The requirement for a functional interaction between the gB and gH/gL ectodomains is consistent with the observation that a purified form of gH/gL ectodomain can to promote a low level of fusion when added exogenously to cells expressing gB and gD (54). Indeed, our previous study identified a potential gB-interaction site in the membrane-proximal domain of the gH ectodomain using species-specific selective pressure (20). Similarly, chimeric constructs of HSV-1 gH and pseudorabies gH also mapped a species-specific gB functional interaction to the membrane-proximal domain of the gH/gL ectodomain (55).

Based on structural analysis of membrane-anchored prefusion gB, a recent study proposed that dissociation of the gB fusion loops from the MPR might trigger gB refolding (14). An interaction with gH/gL at this site in the gB ectodomain could contribute to fusion promotion. The study also identified a nanobody that inhibits gB refolding by binding to prefusion gB between adjacent protomers, contacting domains I, III, and IV. gH/gL interactions that disrupt this gB ectodomain site also might promote fusion.

Importantly, interactions between the gH/gL and gB ectodomains are insufficient to drive full fusion activity. Mutations in the gH TMD or CTD can inhibit or abolish fusion without affecting gH folding and expression (42, 56–60). Conversely, mutations or truncations in the gB CTD can enhance fusion (31, 34, 37, 39, 52). The gB CTD has been proposed to function as a “clamp” that stabilizes gB and regulates its activity by maintaining gB in an inactive conformation until it is triggered (17). The gH CTD has been proposed to act as a “wedge” that disrupts the gB CTD clamp to promote fusion during entry (17). Modeling and mutagenesis of the gB and gH CTD suggests that gH-V831 could interact with a pocket in the gB CTD (45). This site is consistent with our recent findings that identified a potential functional interaction site in the gH CTD using selective pressure with an impaired gB mutant (20).

In support of a model that the gB CTD regulates fusion, our results suggest that the SaHV-1 gB CTD is less stable than the HSV-1 gB CTD. The addition of the HSV-1 CTD to SaHV-1 gB (S1 and S2) enhanced fusion when co-expressed with gD and gH/gL from either species (Fig. 3, 5). Conversely, the addition of the SaHV-1 CTD to HSV-1 gB dampened fusion when the co-expressed with gD and gH/gL from either species. An alignment of the HSV-1 and SaHV-1 gB CTD shows uneven sequence conservation, with greater divergence towards the C-terminal end, especially after HSV-1 residue 884 (Fig. 1C and 1D). Truncations in this region, such as gB876t (Fig. 5), are hyperfusogenic, suggesting that these residues contribute to formation of the CTD clamp structure. The C-terminal residues of SaHV-1 gB may form a tighter clamp than those of HSV-1 gB. Given that WT SaHV-1 gB mediates similar levels of fusion in the cell-cell fusion assay to WT HSV-1 gB, the increased stability of the SaHV-1 gB CTD may be balanced by the SaHV-1 gB ectodomain having less stability. Nevertheless, the importance of homotypic ectodomain indicates that gH/gL interactions with the gB CTD alone are insufficient to trigger fusion.

Because the HSV-1 gB CTD was hyperfusogenic compared to the SaHV-1 gB CTD, we were unable to identify a species-specific functional interaction within the gB CTD. The S1 and S2 chimeras were hyperfusogenic when the target cells expressed nectin-1, HVEM, or CD155 (Fig. 4), indicating that the hyperfusogenicity was not receptor-specific. Although the hyperfusogenic chimeras S1 and S2 appear less stable than WT gB, they retained the gD and gH/gL requirement for triggering (Fig. 6), suggesting that these chimeras still require gH/gL to interact with both the gB ectodomain and CTD.

**Figure 6.**
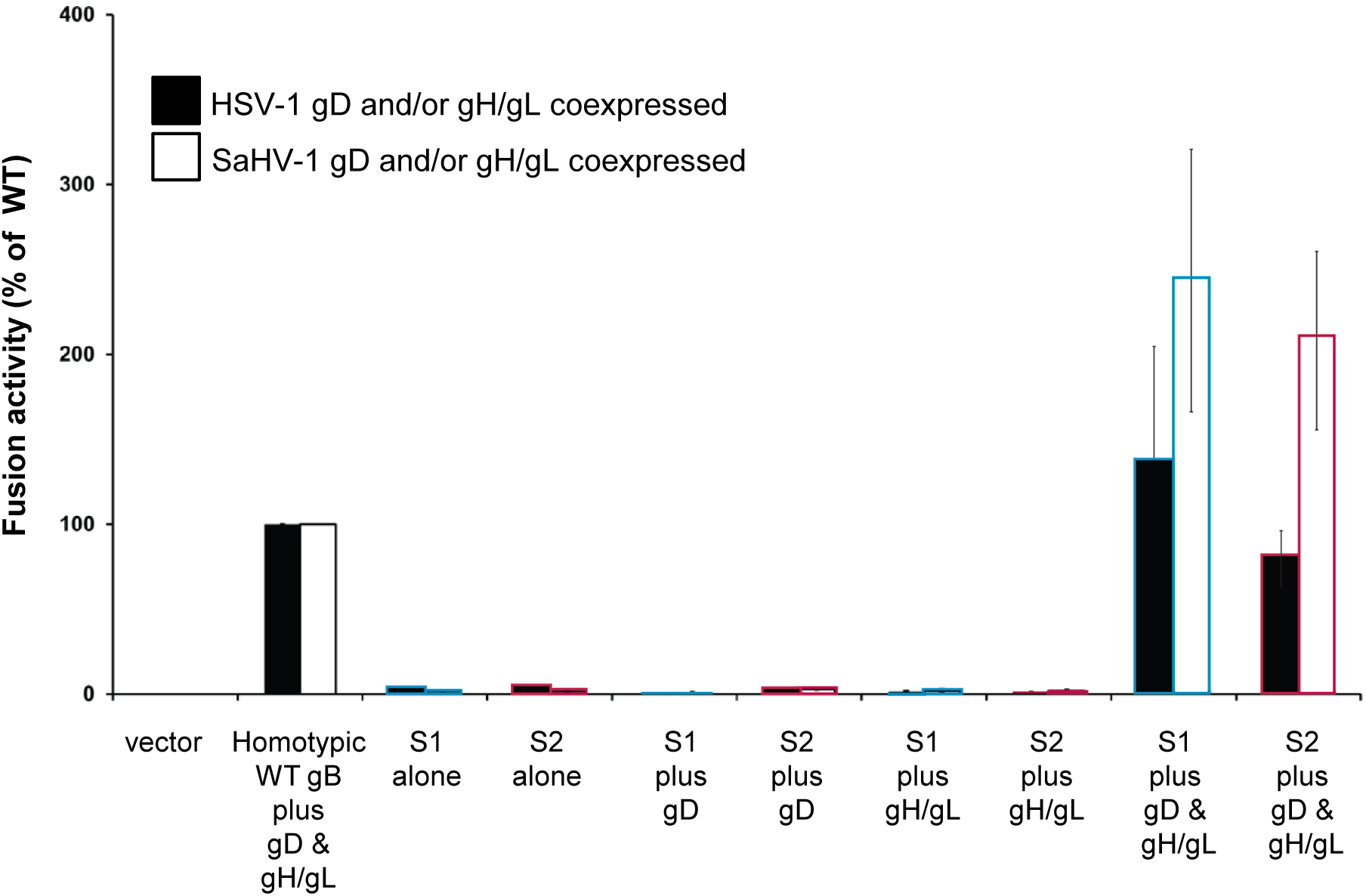
Hyperfusogenic gB chimeras retain dependence on gD and gH/gL. Effector CHO-K1 cells were transfected with plasmids encoding a version of gB (WT, S1 or S2 chimera, or empty vector), gD and/or gH and gL from either HSV-1 (filled bars) or SaHV-1 (open bars), and T7 polymerase. Target cells were transfected with plasmids carrying the luciferase gene under the control of the T7 promoter and nectin-1. Cells were cocultured and luciferase activity was measured. Data are shown as a percentage of WT HSV-1 (filled bars) or WT SaHV-1 (open bars) fusion, following subtraction of vector-only background signal. Means and standard deviations of results of three independent experiments are shown.

Together, these findings refine the current model of herpesvirus-induced membrane fusion by demonstrating that both the ectodomain and cytoplasmic tail domain (CTD) of the conserved viral fusion protein gB participate in regulating fusion. Species-specific functional interactions mapped to the gB ectodomain, indicating that proposed gH-gB CTD interactions alone are not sufficient to trigger fusion. The observation that the HSV-1 gB CTD is less stable than the SaHV-1 gB CTD supports a model that the C-terminal gB residues contributes to a clamp function of the CTD. The robust fusion activity of WT SaHV-1 gB, despite its stable CTD, suggests that herpesviruses may balance stability between the gB ectodomain and CTD, thereby maintaining optimal fusion efficiency during entry.

## ACKNOWLEDGMENTS

We thank Dr. Eckard Wimmer for providing the CD155 expression plasmid. We thank N. Susmarski and Ashley Jiang for timely and excellent technical assistance and members of the Longnecker Laboratory for their help in these studies. Sequencing services were performed at the Northwestern University Genomics Core Facility. R.L. is the Dan and Bertha Spear Research Professor in Microbiology-Immunology. Research reported in this publication was supported by the National Institute of Allergy and Infectious Disease (NIAID) of the National Institutes of Health under grant number AI148478. The content is solely the responsibility of the authors and does not necessarily represent the official views of the National Institutes of Health. This manuscript is the result of funding in whole by the National Institutes of Health (NIH). It is subject to the NIH Public Access Policy. Through acceptance of this federal funding, NIH has been given a right to make this manuscript publicly available in PubMed Central upon the Official Date of Publication, as defined by NIH.

## Data Availability Statement

All data and reagents are available upon request, please contact the corresponding author directly for reuse.

## REFERENCES

1. Connolly SA, Jardetzky TS, Longnecker R. 2021. The structural basis of herpesvirus entry. Nature Reviews Microbiology 19:110–121.

2. Gonzalez-Del Pino GL, Heldwein EE. 2022. Well Put Together-A Guide to Accessorizing with the Herpesvirus gH/gL Complexes. Viruses 14.

3. Connolly SA, Jackson JO, Jardetzky TS, Longnecker R. 2011. Fusing structure and function: a structural view of the herpesvirus entry machinery. Nat Rev Microbiol 9:369–81.

4. Eisenberg RJ, Atanasiu D, Cairns TM, Gallagher JR, Krummenacher C, Cohen GH. 2012. Herpes virus fusion and entry: a story with many characters. Viruses 4:800–32.

5. Stampfer SD, Heldwein EE. 2013. Stuck in the middle: structural insights into the role of the gH/gL heterodimer in herpesvirus entry. Curr Opin Virol 3:13–9.

6. Weed DJ, Nicola AV. 2017. Herpes simplex virus Membrane Fusion. Adv Anat Embryol Cell Biol 223:29–47.

7. Satoh T, Arii J, Suenaga T, Wang J, Kogure A, Uehori J, Arase N, Shiratori I, Tanaka S, Kawaguchi Y, Spear PG, Lanier LL, Arase H. 2008. PILRalpha is a herpes simplex virus-1 entry coreceptor that associates with glycoprotein B. Cell 132:935–44.

8. Arii J, Goto H, Suenaga T, Oyama M, Kozuka-Hata H, Imai T, Minowa A, Akashi H, Arase H, Kawaoka Y, Kawaguchi Y. 2010. Non-muscle myosin IIA is a functional entry receptor for herpes simplex virus-1. Nature 467:859–62.

9. Suenaga T, Satoh T, Somboonthum P, Kawaguchi Y, Mori Y, Arase H. 2010. Myelin-associated glycoprotein mediates membrane fusion and entry of neurotropic herpesviruses. Proc Natl Acad Sci U S A 107:866–71.

10. Heldwein EE, Lou H, Bender FC, Cohen GH, Eisenberg RJ, Harrison SC. 2006. Crystal structure of glycoprotein B from herpes simplex virus 1. Science 313:217–220.

11. Backovic M, Jardetzky TS. 2009. Class III viral membrane fusion proteins. Curr Opin Struct Biol 19:189–96.

12. Cairns TM, Connolly SA. 2021. Entry of Alphaherpesviruses. Curr Issues Mol Biol 41:63–124.

13. Ito F, Zhen J, Xie G, Huang H, Silva JC, Wu TT, Zhou ZH. 2025. Structure of the Kaposi’s sarcoma-associated herpesvirus gB in post-fusion conformation. J Virol 99:e0153324.

14. Vollmer B, Ebel H, Rees R, Nentwig J, Mulvaney T, Schunemann J, Krull J, Topf M, Gorlich D, Grunewald K. 2025. A nanobody specific to prefusion glycoprotein B neutralizes HSV-1 and HSV-2. Nature 646:433–441.

15. Liu Y, Heim KP, Che Y, Chi X, Qiu X, Han S, Dormitzer PR, Yang X. 2021. Prefusion structure of human cytomegalovirus glycoprotein B and structural basis for membrane fusion. Sci Adv 7.

16. Sponholtz MR, Byrne PO, Lee AG, Ramamohan AR, Goldsmith JA, McCool RS, Zhou L, Johnson NV, Hsieh CL, Connors M, Karthigeyan KP, Crooks CM, Fuller AS, Campbell JD, Permar SR, Maynard JA, Yu D, Bottomley MJ, McLellan JS. 2024. Structure-based design of a soluble human cytomegalovirus glycoprotein B antigen stabilized in a prefusion-like conformation. Proc Natl Acad Sci U S A 121:e2404250121.

17. Cooper RS, Georgieva ER, Borbat PP, Freed JH, Heldwein EE. 2018. Structural basis for membrane anchoring and fusion regulation of the herpes simplex virus fusogen gB. Nat Struct Mol Biol 25:416–424.

18. Fan Q, Longnecker R, Connolly SA. 2014. Substitution of herpes simplex virus 1 entry glycoproteins with those of saimiriine herpesvirus 1 reveals a gD-gH/gL functional interaction and a region within the gD profusion domain that is critical for fusion. Journal of virology 88:6470–82.

19. Fan Q, Longnecker R, Connolly SA. 2015. A Functional Interaction between Herpes Simplex Virus 1 Glycoprotein gH/gL Domains I and II and gD Is Defined by Using Alphaherpesvirus gH and gL Chimeras. Journal of Virology 89:7159–7169.

20. Fan Q, Hippler DP, Yang Y, Longnecker R, Connolly SA. 2023. Multiple Sites on Glycoprotein H (gH) Functionally Interact with the gB Fusion Protein to Promote Fusion during Herpes Simplex Virus (HSV) Entry. mBio 14:e0336822.

21. Miller CG, Krummenacher C, Eisenberg RJ, Cohen GH, Fraser NW. 2001. Development of a syngenic murine B16 cell line-derived melanoma susceptible to destruction by neuroattenuated HSV-1. Mol Ther 3:160–168.

22. Herold BC, Visalli RJ, Susmarski N, Brandt CR, Spear PG. 1994. Glycoprotein C-independent binding of herpes simplex virus to cells requires of cell surface heparan sulphate and glycoprotein B. JGenVirol 75:1211–1222.

23. Pertel P, Fridberg A, Parish ML, Spear PG. 2001. Cell fusion induced by herpes simplex virus glycoproteins gB, gD, and gH-gL requires a gD receptor but not necessarily heparan sulfate. Virology 279:313–324.

24. Geraghty RJ, Krummenacher C, Cohen GH, Eisenberg RJ, Spear PG. 1998. Entry of alphaherpesviruses mediated by poliovirus receptor-related protein 1 and poliovirus receptor. Science 280:1618–1620.

25. Montgomery RI, Warner MS, Lum BJ, Spear PG. 1996. Herpes simplex virus-1 entry into cells mediated by a novel member of the TNF/NGF receptor family. Cell 87:427–436.

26. Lin E, Spear PG. 2007. Random linker-insertion mutagenesis to identify functional domains of herpes simplex virus type 1 glycoprotein B. Proc Natl Acad Sci USA 104:13140–13145.

27. Connolly SA, Landsburg DJ, Carfi A, Whitbeck JC, Zuo Y, Wiley DC, Cohen GH, Eisenberg RJ. 2005. Potential nectin-1 binding site on herpes simplex virus glycoprotein D. J Virol 79:1282–1295.

28. Krummenacher C, Rux AH, Whitbeck JC, Ponce-de-Leon M, Lou H, Baribaud I, Hou W, Zou C, Geraghty RJ, Spear PG, Eisenberg RJ, Cohen GH. 1999. The first immunoglobulin-like domain of HveC is sufficient to bind herpes simplex virus gD with full affinity, while the third domain is involved in oligomerization of HveC. J Virol 73:8127–37.

29. Fan Q, Amen M, Harden M, Severini A, Griffiths A, Longnecker R. 2012. Herpes B virus utilizes human nectin-1 but not HVEM nor PILRalpha for cell-cell fusion and virus entry. J Virol doi:10.1128/JVI.00041-12.

30. Atanasiu D, Whitbeck JC, de Leon MP, Lou H, Hannah BP, Cohen GH, Eisenberg RJ. 2010. Bimolecular complementation defines functional regions of Herpes simplex virus gB that are involved with gH/gL as a necessary step leading to cell fusion. J Virol 84:3825–34.

31. Chowdary TK, Cairns TM, Atanasiu D, Cohen GH, Eisenberg RJ, Heldwein EE. 2010. Crystal structure of the conserved herpesvirus fusion regulator complex gH-gL. Nat Struct Mol Biol 17:882–8.

32. Cai WZ, Person S, Debroy C, Gu BH. 1988. Functional regions and structural features of the gB glycoprotein of herpes simplex virus type 1. An analysis of linker insertion mutants. JMolBiol 201:575–588.

33. Wanas E, Efler S, Ghosh K, Ghosh HP. 1999. Mutations in the conserved carboxy-terminal hydrophobic region of glycoprotein gB affect infectivity of herpes simplex virus. J Gen Virol 80 (Pt 12):3189–3198.

34. Baghian A, Huang L, Newman S, Jayachandra S, Kousoulas KG. 1993. Truncation of the carboxy-terminal 28 amino acids of glycoprotein B specified by herpes simplex virus type 1 mutant amb1511-7 causes extensive cell fusion. JVirol 67:2396–2401.

35. Diakidi-Kosta A, Michailidou G, Kontogounis G, Sivropoulou A, Arsenakis M. 2003. A single amino acid substitution in the cytoplasmic tail of the glycoprotein B of herpes simplex virus 1 affects both syncytium formation and binding to intracellular heparan sulfate. Virus Research 93:99–108.

36. Engel JP, Boyer EP, Goodman JL. 1993. Two novel single amino acid syncytial mutations in the carboxy terminus of glycoprotein B of herpes simplex virus type 1 confer a unique pathogenic phenotype. Virology 192:112–120.

37. Fan Z, Grantham ML, Smith MS, Anderson ES, Cardelli JA, Muggeridge MI. 2002. Truncation of herpes simplex virus type 2 glycoprotein B increases its cell surface expression and activity in cell-cell fusion, but these properties are unrelated. J Virol 76:9271–9283.

38. Foster TP, Melancon JM, Kousoulas KG. 2001. An α-helical domain within the carboxyl terminus of herpes simplex virus type1 (HSV-1) glycoprotein B (gB) is associated with cell fusion and resistance to heparin inhibition of cell fusion. Virology 287:18–29.

39. Gage PJ, Levine M, Glorioso JC. 1993. Syncytium-inducing mutations localize to two discrete regions within the cytoplasmic domain of herpes simplex virus type 1 glycoprotein B. JVirol 67:2191–2201.

40. Heineman TC, Hall SL. 2002. Role of the varicella-zoster virus gB cytoplasmic domain in gB transport and viral egress. J Virol 76:591–9.

41. Silverman JL, Greene NG, King DS, Heldwein EE. 2012. Membrane requirement for folding of the herpes simplex virus 1 gB cytodomain suggests a unique mechanism of fusion regulation. J Virol 86:8171–84.

42. Rogalin HB, Heldwein EE. 2015. Interplay between the Herpes Simplex Virus 1 gB Cytodomain and the gH Cytotail during Cell-Cell Fusion. J Virol 89:12262–72.

43. Ruel N, Zago A, Spear PG. 2006. Alanine substitution of conserved residues in the cytoplasmic tail of herpes simplex virus gB can enhance or abolish cell fusion activity and viral entry. Virology 346:229–237.

44. Walev I, Lingen M, Lazzaro M, Weise K, Falke D. 1994. Cyclosporin A resistance of herpes simplex virus-induced “fusion from within” as a phenotypical marker of mutations in the syn 3 locus of the glycoprotein B gene. Virus Genes 8:83–86.

45. Pataki Z, Sanders EK, Heldwein EE. 2022. A surface pocket in the cytoplasmic domain of the herpes simplex virus fusogen gB controls membrane fusion. PLoS Pathog 18:e1010435.

46. Miller CG, Krummenacher C, Eisenberg RJ, Cohen GH, Fraser NW. 2001. Development of a syngenic murine B16 cell line-derived melanoma susceptible to destruction by neuroattenuated HSV-1. Mol Ther 3:160–8.

47. Cairns TM, Atanasiu D, Saw WT, Lou H, Whitbeck JC, Ditto NT, Bruun B, Browne H, Bennett L, Wu C, Krummenacher C, Brooks BD, Eisenberg RJ, Cohen GH. 2020. Localization of the Interaction Site of Herpes Simplex Virus Glycoprotein D (gD) on the Membrane Fusion Regulator, gH/gL. J Virol 94.

48. Cairns TM, Ditto NT, Atanasiu D, Lou H, Brooks BD, Saw WT, Eisenberg RJ, Cohen GH. 2019. Surface Plasmon Resonance Reveals Direct Binding of Herpes Simplex Virus Glycoproteins gH/gL to gD and Locates a gH/gL Binding Site on gD. J Virol 93.

49. Atanasiu D, Saw WT, Cairns TM, Eisenberg RJ, Cohen GH. 2021. Using Split Luciferase Assay and anti-HSV Glycoprotein Monoclonal Antibodies to Predict a Functional Binding Site Between gD and gH/gL. J Virol doi:10.1128/JVI.00053-21.

50. Cairns TM, Whitbeck JC, Lou H, Heldwein EE, Chowdary TK, Eisenberg RJ, Cohen GH. 2011. Capturing the herpes simplex virus core fusion complex (gB-gH/gL) in an acidic environment. J Virol 85:6175–84.

51. Nicola AV. 2016. Herpesvirus Entry into Host Cells Mediated by Endosomal Low pH. Traffic 17:965–975.

52. Gianopulos KA, Makio AO, Pritchard SM, Cunha CW, Hull MA, Nicola AV. 2024. Herpes Simplex Virus 1 Glycoprotein B from a Hyperfusogenic Virus Mediates Enhanced Cell-Cell Fusion. Viruses 16.

53. Atanasiu D, Whitbeck JC, Cairns TM, Reilly B, Cohen GH, Eisenberg RJ. 2007. Bimolecular complementation reveals that glycoproteins gB and gH/gL of herpes simplex virus interact with each other during cell fusion. Proc Natl Acad Sci USA 104:18718–18723.

54. Atanasiu D, Saw WT, Cohen GH, Eisenberg RJ. 2010. Cascade of events governing cell-cell fusion induced by herpes simplex virus glycoproteins gD, gH/gL, and gB. J Virol 84:12292–9.

55. Bohm SW, Backovic M, Klupp BG, Rey FA, Mettenleiter TC, Fuchs W. 2016. Functional Characterization of Glycoprotein H Chimeras Composed of Conserved Domains of the Pseudorabies Virus and Herpes Simplex Virus 1 Homologs. J Virol 90:421–32.

56. Browne HM, Bruun BC, Minson AC. 1996. Characterization of herpes simplex virus type 1 recombinants with mutations in the cytoplasmic tail of glycoprotein H. JGenVirol 77:2569–73.

57. Harman A, Browne H, Minson T. 2002. The Transmembrane Domain and Cytoplasmic Tail of Herpes Simplex Virus Type 1 Glycoprotein H Play a Role in Membrane Fusion. J Virol 76:10708–10716.

58. Jackson JO, Lin E, Spear PG, Longnecker R. 2010. Insertion mutations in herpes simplex virus 1 glycoprotein H reduce cell surface expression, slow the rate of cell fusion, or abrogate functions in cell fusion and viral entry. J Virol 84:2038–46.

59. Silverman JL, Heldwein EE. 2013. Mutations in the cytoplasmic tail of herpes simplex virus 1 gH reduce the fusogenicity of gB in transfected cells. J Virol 87:10139–47.

60. Wilson DW, Davis-Poynter N, Minson AC. 1994. Mutations in the cytoplasmic tail of herpes simplex virus glycoprotein H suppress cell fusion by a syncytial strain. JVirol 68:6985–6993.

